# Provisioning polyethylene glycol (PEG) to large herbivores in nutrient poor savannas can break food limitation

**DOI:** 10.1101/2024.01.20.576483

**Authors:** Bradley Schroder, Frank Van Langevelde, Nicola-Anne Hawkins Schroder, Herbert H. T. Prins

**Affiliations:** Wildlife Ecology and Conservation Group, Wageningen University, Droevendaalsesteeg 3a, 6708 PB Wageningen, the Netherlands; No 13 Upper Waterkloof, 173 Regulus Street, Waterkloof, 0181, Pretoria, South Africa; Animal Sciences Group, Wageningen University, De Elst 1, 6708 WD Wageningen, the Netherlands

**Author notes:** Corresponding author: Dr Bradley Schroder.

**Keywords:** diet selection, food quality, game lick blocks, Polyethylene glycol (PEG), secondary compounds

## Abstract

Reproduction and survival of herbivores in nutrient poor savannas is low due to low nutrient and energy availability, partly caused by high levels of tannins. Polyethylene glycol (PEG) increases the availability of proteins for herbivores by binding tannins. The effect of PEG on the diet of free-roaming herbivores has not been tested. Our hypothesis was deploying lick blocks with the addition of PEG in a nutrient poor savanna, will result in a broadening of the diet of free-roaming herbivores with higher percentages of browse species and higher utilisation per browse species, with higher tannin levels. We further hypothesised that the mineral content in the faeces, once exposed to PEG would increase. We collected faecal samples of five herbivore species with various feeding methods (grazers, browsers or mixed feeders). The study used a Before-After-Control-Impact (BACI) design. The results show that the addition of PEG promotes a change in the browse dietary choices of four of the five herbivore species, and are expressed as a broader choice of diet, coupled with higher numbers of browse species with low edibility and higher tannin levels. The addition of PEG had no noteworthy effect on the concentration of minerals found in the faeces.

## Introduction

Nutrient poor savannas are dominated by broadleaved trees, having leaves with low nutritional value due to high levels of secondary compounds, such as condensed tannin and other carbon-based secondary metabolites (Parker 2004; Mucina and Rutherford 2006; Hattas 2014). The secondary compounds are thought to be the plant’s evolutionary defence response to herbivore utilisation (Cooper and Owen-Smith 1985; Hattas 2014; Scogings et al. 2014). Plants with high levels of secondary compounds can have relatively high nutrient concentrations in their leaves (Tomlinson et al. 2016). It is well documented that condensed tannins occur in a wide range of plant species in nutrient poor savannas (Salawu et al. 1997) and that tannins bind with proteins and often suppress the food intake and digestion by mammalian herbivores (McLeod 1974; Cooper and Owen-Smith 1985; Smit 2013; Scogings et al. 2014; Mkhize et al. 2015). Browsers and mixed feeders’ foraging preferences are strongly shaped by secondary compounds (Scogings et al. 2014).

Polyethylene glycol (PEG) (with commercial names such as Browse Plus or Movicol) is an inert and unabsorbed molecule, which binds with tannins to form a stable complex, preventing the binding between tannins and proteins (Badran and Jones 1965; Makkar et al. 1995; Majuva-Masafu and Linington 2006) increasing the availability of proteins. PEG may increase the availability of macronutrients for herbivores and decrease malaise (Provenza et al. 2000). To date, several studies have shown the effect of chemical defences and PEG on the diet of mammalian herbivores, such as goats (*Capra aegagrus hircus*), sheep (*Ovis aries*) and cattle (B*os taurus africanus*) (Salawu et al. 1997; Decandia et al. 2000; Moujahed et al. 2000; Landau et al. 2003; Mkhize et al. 2015). Experiments have been undertaken on penned sheep and goats, with results showing that tannins not only act as digestibility reducers, but also as feeding deterrents that affect intake, feeding behaviour and diet choice of these herbivores (Foley and Hume 1987; Marsh et al. 2003; Jansen et al. 2007; Mkhize et al. 2015; Mkhize et al. 2016). PEG has been shown to reduce these adverse effects of dietary tannins in these domestic animal’s diets (Salawu et al. 1997; Moujahed et al. 2000; Mkhize et al. 2018). However, little is known about the effect of PEG on free-roaming mammalian herbivores in nutrient poor savannas. Free-roaming herbivores encounter plant species that largely differ in concentrations of nutrients and plant secondary compounds and they have the opportunity to select their diet to counteract the differences in forage quality and plant defences (Prins and Van Langevelde 2008). According to Prins and Van Langevelde (2008) and Provenza et al. (2007), the fitness of herbivores increases when they have access to a variety of plant species with different concentrations of nutrients and secondary compounds, than when constrained to a single food source.

In many game reserves in South Africa, especially on nutrient poor soils, nutrient lick blocks (standard game lick blocks) have been used, originally devised for domestic animals and now designed to supplement the diet of wild animals (Langman 1978; Makkar et al. 2007; SAFARI Feeds 2019; WES Feeds 2019) (see Supporting Information A for the mineral composition of the standard game lick block in South Africa). Putman and Staines (2004) reviewed a number of studies that show experimentally that supplementary feeding free-roaming herbivores with game lick blocks resulted in an improvement of the wild animal’s health and reproductive life cycle. With the addition of PEG to game lick blocks, we expect that large herbivores are able to access previously unusable proteins, indigestible fibres and other sources of energy, locked in plants which would otherwise be unpalatable. This allows the herbivores to include additional plant species in their diet that have levels of condensed tannins that are so high that they would not normally be consumed by herbivores without PEG supplementation, especially during the dry winter months when good quality food is scarce. Our study analyses the change in the diet of free-roaming mammalian herbivore species in response to game lick blocks with and without the addition of PEG. Our hypothesis is that deploying game lick blocks with PEG compared to the standard game lick blocks in a nutrient poor savanna will result in a broadening of the diet of herbivores with a higher percentages of browse species and higher amounts of utilisation per browse species that have low palatability and generally have higher levels of condensed tannins. We further hypothesised that the mineral content within the faeces exposed to the game lick blocks with PEG would have an increase in mineral concentrations. To test for differences in the quality of the diet after exposure to the game lick blocks, the minerals were compared in the faeces of mammalian herbivores exposed to standard game lick blocks versus those exposed to the game lick blocks with the addition of PEG. An increase in the concentration of minerals in the faeces would potentially show an increase in the quality of minerals obtained from the increase in diet range and quantity.

## Material and methods

### Study area

The study was conducted in the Welgevonden Game Reserve (348 km²), situated on the Waterberg Plateau in South Africa (24°10’S; 27°45’E to 24°25’S; 27°56’E), over a period of three years (2016 – 2018). The area is classified as warm and temperate, with summer rainfall, having distinct wet and dry seasons that stretch from October to March and April to September respectively, with a mean annual rainfall off 665 mm. The mean annual maximum temperature is 27.4°C (reaching 40°C) and the mean annual minimum temperature is 14.5°C (reaching -4°C). Welgevonden Game Reserve is situated in two biomes, the Savanna Biome and Grassland Biome, and falls mainly within the Waterberg Mountain Bushveld vegetation type (Mucina and Rutherford 2006). The area is characterised primarily by the soil types dystrophic to mesotrophic yellow-brown apedal coarse sands (Parker 2004), with ferruginous soils with a low pH. Accordingly, the vegetation type is dominated by nutritionally poor broadleaved savanna. Mucina and Rutherford (2006) classify this area as a nutrient poor savanna ecosystem (locally termed ‘sour veld’, a.k.a. ‘dystrophic savanna’). The study area comprises mountainous terrain that is dissected by deep valleys, with occasional old agricultural lands. Flat plateaus characterise most of the hilltops, and the elevation varies from 1050 m to 1800 m above sea level. The previous land use included agriculture, cattle ranching and hunting, with the area now only being utilised for conservation and eco-tourism. Sixty-three species of free-roaming mammals have been identified in the study area, including various antelope species, mega-herbivores and predators.

### Methods

We studied five mammalian herbivore species to establish the utilisation of PEG and the effect on herbivore diets: one mixed feeder, *viz*., impala (*Aepyceros melampus*), two browsers, *viz.,* greater kudu (*Tragelaphus strepsiceros*) and the common eland (*Taurotragus oryx*), and two grazers, *viz.,* Burchell’s zebra (*Equus quagga burchellii*) and the blue wildebeest (*Connochaetes taurinus*). To test the effect of the standard game lick block and the PEG game lick block on the diet of these herbivore species a Before-After-Control-Impact (BACI) design was used.

The topography of Welgevonden Game Reserve is determined by its underlying geology (*viz*., an uplifted sandstone plateau crisscrossed by rifts) which creates valleys flanked by steep hill slopes. Due to the steep topography the normal movement patterns of herbivore species is primarily along the valleys (Mr. J Swart 2017, personal communication) and rarely between them. Six sites were identified within six valleys, which were roughly 4 – 5 kms apart, to prevent individual animals visiting more than one site during the study. A ‘transect’ of 12 game lick blocks were aligned at 25m intervals running centrally through each valley (Figure 1, shows the layout design of the game lick blocks in the study area). The transect layout maximised the chance that the animals would eat and defecate in the same area ensuring the faecal samples obtained for analysis were from animals which had consumed either the standard or PEG game lick block, post deployment of the lick blocks.

**Figure 1.**
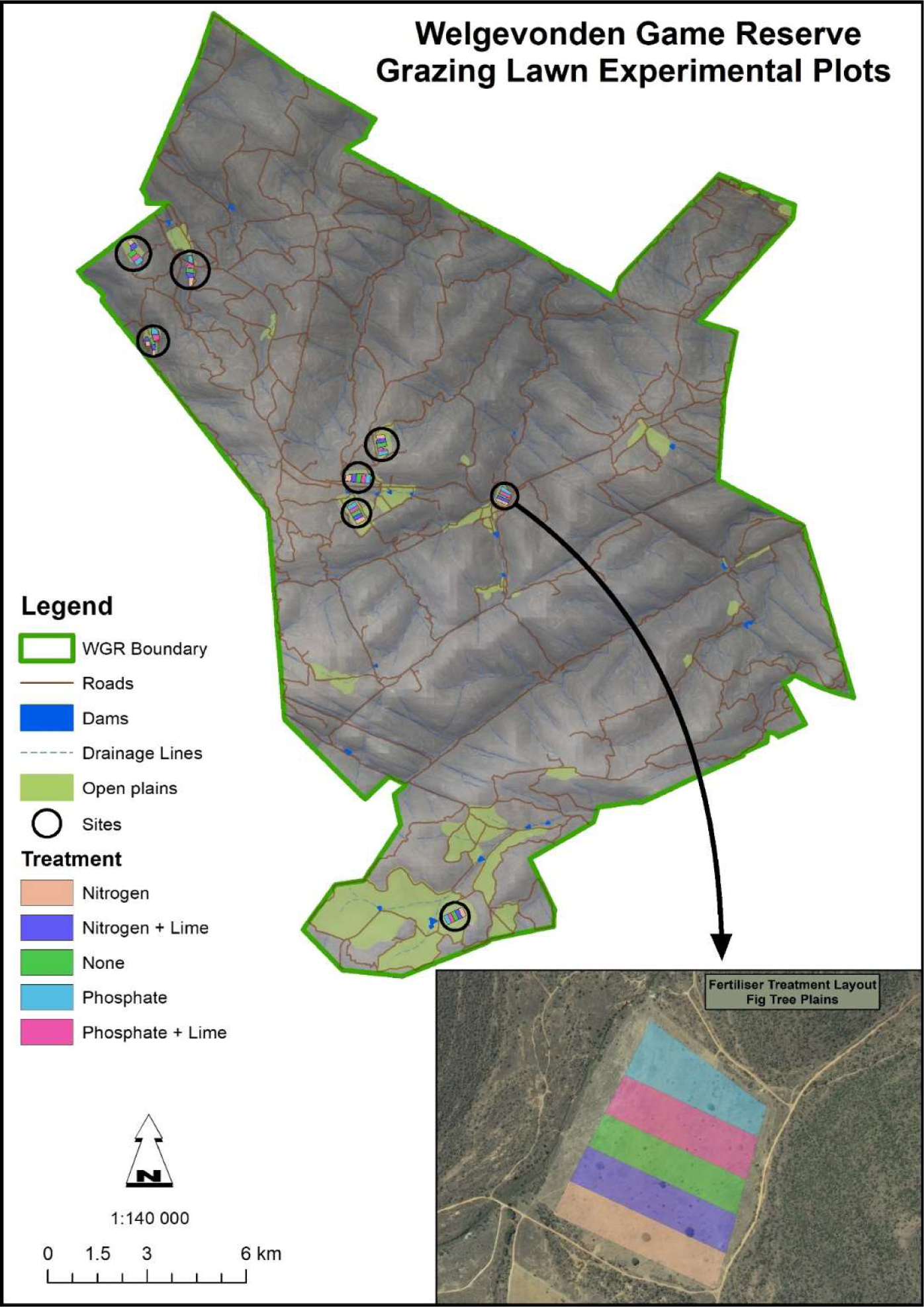
Map of the Welgevonden Game Reserve showing the grazing lawn experimental sites and layout of treatment plots. WGR = Welgevonden Game Reserve

Prior to the game lick blocks being deployed faecal samples of the five-herbivore species were collected from the six predetermined sites (‘Before Assessment’ analysis). Samples were collected from the area 25m prior to where the first lick block would be deployed and to 25m after where the last lick block would be deployed and at a diameter of 25m either side of the transect. Two types of game lick blocks (each with an individual mass of 25kg) were then deployed in the study area at the six sites (three sites per game lick block type). The first type was the standard game lick block (of which the nutrient supplements are given in Supporting Information A), the second was the standard game lick blocks with the addition of 35g of PEG per 25kg block (WES Feeds 2019, personal communication as per our request). The game lick blocks were placed along a transect within each site for consumption during the dry months of the year (May through October). The game lick blocks were replaced every two to three weeks until the end of October. A second collection of faecal samples were collected in early November (‘After Assessment’ analysis). All the herbivore species could voluntarily choose to consume or not to consume the game lick blocks. As dictated by standard practise, the game lick blocks were not provided during the wet season (November through May), as they can negatively affect herbivores.

During this experiment, the health and welfare of the free-roaming animals was observed and checked regularly and at no stage during this experiment were any animals harmed.

### Data Collection

The experiment was repeated over a three-year period (2016 to 2018). The faecal sample collection prior to the game lick block application was undertaken in April of each year. The game lick blocks were placed in transects at the beginning of May and replaced every two to three weeks until the end of October, which is a period of six months each year. The faecal sample collection after the game lick block applications was done in early November of each year. A minimum of five individual faecal samples per animal species were collected (each sample was no less than five pellets), if there were less than five samples per animal species to collect, that species faeces were not included in the site data analysis for that month. The sample was estimated to be no older than five days based on water retention and colour. Over the three-year period 655 individual faecal samples (different animal droppings from each of the five species) were collected from the five species. The faecal samples collected per species were represented as follows, impala (145), greater kudu (119), common eland (80), Burchell’s zebra (156) and blue wildebeest (155).

Each faeces sample was analysed for mineral concentrations and the presence of individual grass and woody plant species. The mineral concentrations of the faeces were analysed after the six-month period of game block deployment/application to compare the mineral concentration between herbivores exposed to the standard verses PEG game lick blocks. The minerals analysed in the faeces have important uses for the five large free-roaming herbivores (Table 1) and were analysed either as a percentage in the case of (N, Ca, Mg, K, S and P) or as mg/kg as with (Na, Fe, Mn, Cu, Zn and B) using the Combustion method (DUMAS test) for N and the Inductively coupled plasma optical emission spectroscopy (ICP-OES) test for the remaining minerals.

**Table 1.** Important use of minerals and elements for free-roaming herbivores (Roosendaal 1992; McDowell & Arthington 2005) which are included in the faeces mineral concentration analysis

| **Nutrient/Element/Mineral** | **Use in free-roaming herbivores** |
| --- | --- |
| Sodium (Na) | Used by animals in extracellular fluids such as blood plasma |
| Nitrogen (N) | A major component of protein in animals and used for growth, reproduction and survival |
| Calcium (Ca) | Mineral found in greatest abundances in animals and used for structural support especially in the bones |
| Magnesium (Mg) | Important mineral assisting with nerve and muscle function, immune system function and bone health |
| Potassium (K) | Necessary for the function of all living cells |
| Sulphur (S) | Important for ruminants for growth and milk production |
| Phosphorus (P) | Essential mineral for cell development and a key component to store energy |
| Iron (Fe) | Used to produce haemoglobin, which delivers oxygen to the blood and carbon dioxide out of the body |
| Manganese (Mn) | Important trace mineral for enzyme activation and stimulates growth |
| Copper (Cu) | Essential trace element required for body, bone and white blood cell function |
| Zinc (Zn) | Used for cellular respiration, cellular use of oxygen and maintenance of cell membranes |
| Boron (B) | Used for immune response, antioxidant, bone metabolism and enhancing animal performance |

The faecal mineral content analyses are useful as short-term indicators of diet selection and nutrient status of free-roaming herbivores (Botha and Stock 2005; Codron et al. 2005).

The presence of individual grass and woody plant species were analysed using microhistological faecal analysis via observation of plant cuticles or epidermal fragments found in the faeces. The cuticle and epidermis of plants have specific structures and patterns so they can be individually identified using a microscope, to the plant species level (De Jong et al. 1995). This analysis allowed us to establish the individual species of browse and graze that the animals had been utilising. 100 fragments per faeces sample were randomly selected and analysed and then used to describe the animal’s diet (Van Lieverloo et al. 2009). The number of individual fragments per species within the faeces sample, could be used as a proxy for the amount of materials per plant species consumed by the animal (De Jong et al. 1995). The Welgevonden Game Reserve plant species collection list was used as the reference to authenticate the species identified within the faeces samples. Both the woody plant species and grass species identified in the faeces were ranked according to their estimated palatability (Supporting Information B and C). Palatability is difficult to define in terms of the biological processes involved in food selection. As commonly used the term implies acceptability but not necessarily desirability (Molyneux and Ralphs 1992). Tree species digestibility ratings were established through the use of the Field Guide to TREES of Southern Africa (Van Wyk and Van Wyk 1997) and through the independent advice of experts in the field (Supporting Information B). Grazing values of the grasses were obtained from the Guide to Grasses of southern Africa (Van Oudtshoorn 2002). The faeces were not analysed for herbs (forbs).

In Supporting Information D and E, the individual species of browse and grass found in the faeces of the five-animal species during the study period are listed, together with the palatability of plant species and grazing ratings of grass species, coupled with the diet change per herbivore species.

### Statistical Analysis

The change in diet was analysed when comparing before versus after deployment of the standard game lick blocks and the PEG augmented game lick blocks. Therefore, comparing plant species composition in the diets using a Principal Component Analysis (PCA). The PCA allowed for the correlation in the change in diet with the quality of the plant species (palatability of the tree species and grazing values for the grass species). The hypothesis was tested using linear mixed models (LMMs) with mineral concentrations of the faeces and the percentage of browse species in the diet as response variables, and animal species, before versus after and PEG-augmented game lick blocks versus standard game lick blocks as fixed factors and year and site as random factors. The differences between the animal species were analysed using Šidák multiple comparisons tests. If the residuals of the Linear mixed models were not normally distributed transformation of the response variable was performed. Analysis for the PCA were performed in Canoco 5.0 (Šmilauer and Lepš 2014) and all other analysis was performed using SPSS v. 23 (IBM SPSS Inc., Chicago, USA).

## Results

### Percentage browse in diet

The addition of PEG to the standard game lick blocks promotes a change in the dietary choices of four of the five herbivore species namely the common eland, greater kudu, Burchell’s zebra and impala. These changes can be expressed as a broader diet choice, either as a higher percentage of browse species or a higher number of individual species utilised with higher concentrations of secondary compounds and low to average palatability in their diet. We found that due to the consumption of the PEG game lick blocks, two of the five herbivore species, impala and Burchell’s zebra, had statistically significant changes in the percentage of browse utilised in their diet. (Figure 2, Table 2). There were smaller changes in increased browse utilisation in the common eland and greater kudu. There was a decrease in browse utilisation by blue wildebeest after the consumption of the PEG game lick block (Table 3).

**Figure 2.**
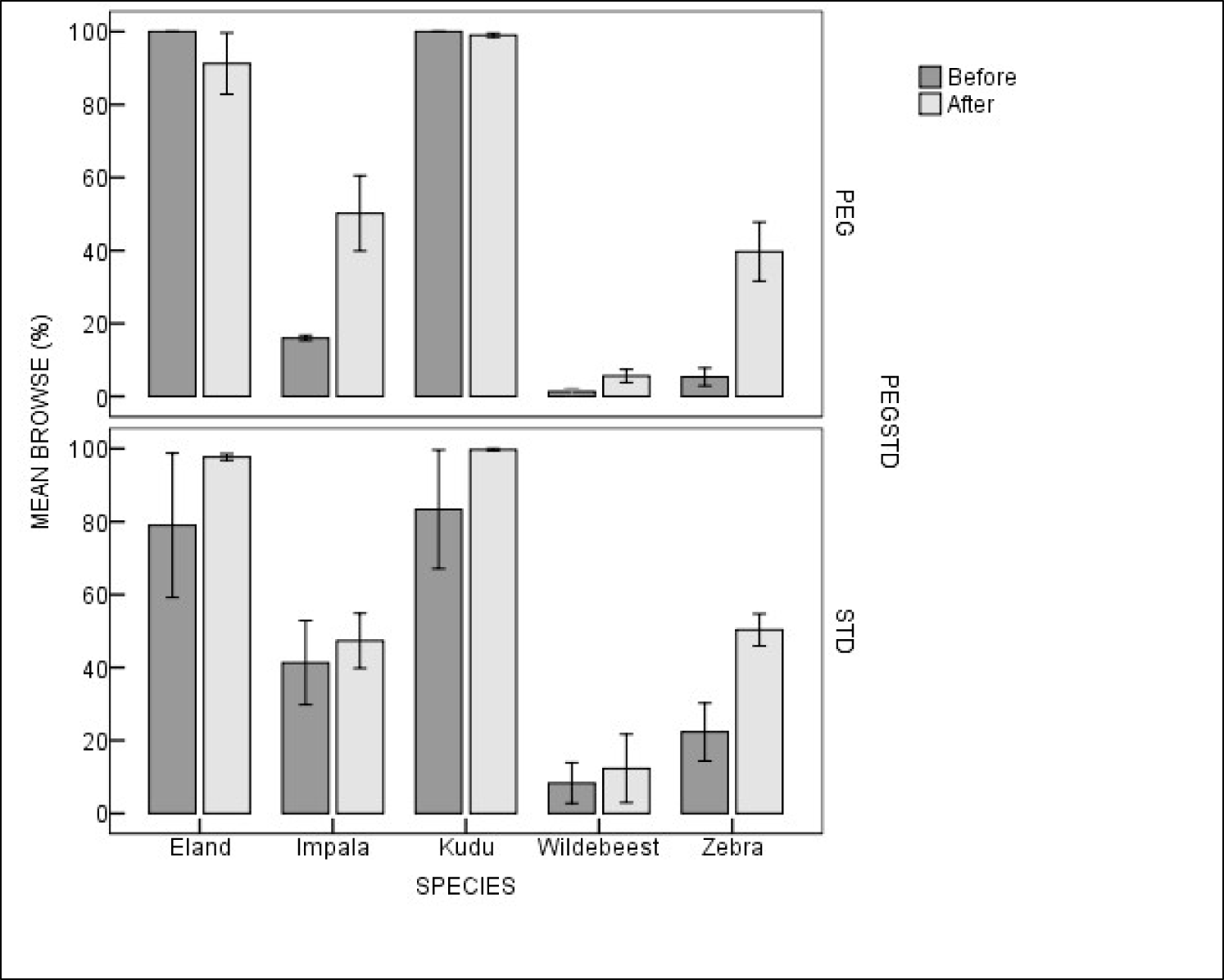
Percentage of total browse species consumption, over a 3-year period, collected in the Welgevonden Game Reserve, South Africa, for each herbivore species (*viz*. impala (*Aepyceros melampus*), greater kudu (*Tragelaphus strepsiceros*), common eland (*Taurotragus oryx*), Burchell’s zebra (*Equus quagga burchellii*) and the blue wildebeest (*Connochaetes taurinus*) per game lick block type. Before and After refers to the percentage of total browse species in the diet taken before or after the utilisation of either the standard game lick blocks (STD) or the game lick block with the addition of Polyethylene glycol (PEG). Error bars represent the standard error of the mean. The stars indicate the statistically significant differences between the STD and PEG game lick blocks per species and per treatment based on Šidák multiple comparisons tests using a linear mixed model (see Table 2 for the statistics)

**Table 2.** Results of the linear mixed model for differences in percentage of browse species in the faeces before and after applying the Standard (STD) and Polyethylene glycol (PEG) game lick blocks (“block type”) for the five herbivore species (*viz*. impala (*Aepyceros melampus*), greater kudu (*Tragelaphus strepsiceros*), common eland (*Taurotragus oryx*), Burchell’s zebra (*Equus quagga burchellii*) and the blue wildebeest (*Connochaetes taurinus*). Year and site were the random factors, estimation methods were REML and the sample size n = 91 (see Figure 3 for the graphs). Akaike Information Criterion (AIC) estimates the prediction error of the amount of information lost in the model and establishes the quality of the model. This data was collected over a three-year period, 2016 – 2018, on the Welgevonden Game Reserve, South Africa. N.S. = not significant at the 5% level

| Herbivore Species |  | Browse |
| --- | --- | --- |
| Herbivore Species | F | 185.6 |
|  | df | 124.7 |
|  | P | <0.001 |
| Block Type | F | 0.08 |
|  | df | 125.8 |
|  | P | N.S. |
| Before - After | F | 5.5 |
|  | df | 126.2 |
|  | P | <0.05 |
| Herbivore species x Block Type | F | 1.2 |
|  | df | 124.5 |
|  | P | N.S. |
| Herbivore species x Before - After | F | 4.0 |
|  | df | 124.6 |
|  | P | <0.01 |
| Before - After x Block Type | F | 0.912 |
|  | df | 126.2 |
|  | P | N.S. |
| Herbivore species x Before - After x Block Type | F | 1.0 |
|  | df | 124.5 |
|  | P | N.S. |
| Year | Wald Z | 0.8 |
|  | P | N.S. |
| Site | Wald Z | 0.4 |
|  | P | N.S. |
| AIC |  | 1126.2 |

**Table 3.** Percentage change in the amount of browse and graze fragments in the diet of each individual free-roaming herbivore species (viz. impala (*Aepyceros melampus*), greater kudu (*Tragelaphus strepsiceros*), common eland (*Taurotragus oryx*), Burchell’s zebra (*Equus quagga burchellii*) and the blue wildebeest (*Connochaetes taurinus*), per game lick block type collected over a three-year period, 2016 – 2018, in the Welgevonden Game Reserve, South Africa

| Species | Lick block type | Browse |  |  |  |  | Graze |  |  |  |  |
| --- | --- | --- | --- | --- | --- | --- | --- | --- | --- | --- | --- |
|  |  | No. Browse species fragments before lick block deployment | No. Browse species fragments after lick block deployment | % Browse species fragments before lick block deployment | % Browse species fragments after lick block deployment | Change in browse utilisation | No. Graze species fragments before lick block deployment | No. Graze species fragments after lick block deployment | % Graze species fragments before lick block deployment | % Graze species fragments after lick block deployment | Change in graze utilisation |
| Burchell's zebra | PEG | 44 | 314 | 8% | 35% | 27% | 519 | 577 | 92% | 65% | -27% |
|  | STD | 141 | 196 | 16% | 32% | 17% | 757 | 412 | 84% | 68% | -17% |
| Blue wildebeest | PEG | 49 | 31 | 10% | 4% | -6% | 447 | 761 | 90% | 96% | 6% |
|  | STD | 59 | 50 | 6% | 8% | 2% | 881 | 550 | 94% | 92% | -2% |
| Greater kudu | PEG | 385 | 779 | 99% | 99% | 0% | 5 | 7 | 1% | 1% | 0% |
|  | STD | 778 | 513 | 88% | 86% | -2% | 104 | 84 | 12% | 14% | 2% |
| Impala | PEG | 106 | 316 | 18% | 45% | 27% | 487 | 381 | 82% | 55% | -27% |
|  | STD | 281 | 220 | 31% | 37% | 5% | 615 | 380 | 69% | 63% | -5% |
| Common eland | PEG | 397 | 560 | 99% | 93% | -6% | 3 | 39 | 1% | 7% | 6% |
|  | STD | 688 | 591 | 99% | 99% | 0% | 9 | 8 | 1% | 1% | 0% |

There were particularly large differences in the number of browse species utilised by the impala and Burchell’s zebra after the additional of PEG in their diet. The number of browse species utilised increased by 181% in impala and 194% in Burchell’s zebra (supporting information D, E and Table 3). There was an increase in the total fragments of browse found in the faeces of Burchell’s zebra, kudu, impala and common eland after the utilization of PEG, compared with the standard lick block. Wildebeest was the only species with a decrease in total fragments of browse after the addition of PEG (Table 3).

For Burchell’s zebra we found a significant change in their diet from grass to browse with the utilisation of PEG, from a diet of 8% browse to 35% browse with an increase from 16 to 31 different browse species, increasing the use of less palatable species (Supporting Information D, E and Table 3). There was an increase in the utilisation of browse after the use of the standard game lick blocks but not nearly as large as that after the addition of PEG.

### Diet composition

The changes in diet composition in terms of the different browse and grass species for the five species of herbivores after the introduction of the standard game lick blocks and PEG game lick blocks are shown in Figure 3, with ‘Panel a’ encompassing the grass species results and ‘Panel b’ the browse species results.

**Figure 3.**
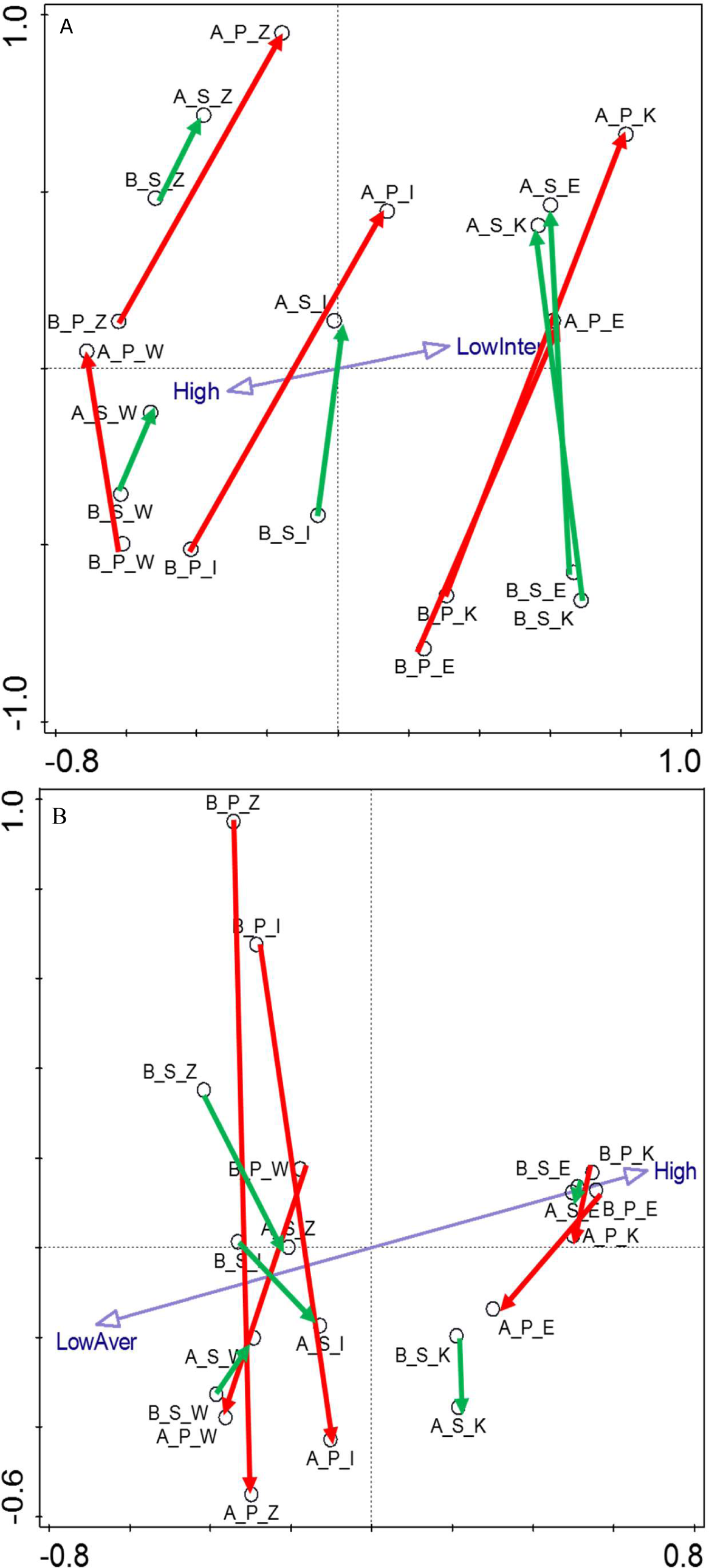
Diagrams of the first two axes of the Principal Components Analysis based on (Panel a) the grass species and (Panel b) the browse species in the diets of the five free-roaming herbivore species (*viz*. impala (*Aepyceros melampus*), greater kudu (*Tragelaphus strepsiceros*), common eland (*Taurotragus oryx*), Burchell’s zebra (*Equus quagga burchellii*) and the blue wildebeest (*Connochaetes taurinus*) before and after the utilization of the Standard (STD) game lick blocks or Polyethylene glycol (PEG) game lick blocks. Every dot represents the combined diet of a herbivore species before or after the addition of the standard game lick blocks or the PEG game lick blocks. The location of these dots is determined by the plant species composition of the diet and the abundance of that plant species that was consumed, so dots close to each other have more similar diets than dots further apart. The red arrows show changes in diet following the application of the PEG game lick blocks, whereas the green arrows refer to the changes in diet after the deployment of STD blocks. The blue arrows show the gradients in the foraging ratings (in panel a; classes low and intermediate are merged and shown as “LowInter”) and palatability (in panel b; classes low and average are merged and shown as “LowAver”) in the diet. Abbreviations: A = After; B = Before; S = STD; P = PEG; E = Eland; I = Impala; K = Kudu; W = Wildebeest and Z = Zebra. The faecal samples were collected over a three-year period, 2016 – 2018, in the Welgevonden Game Reserve, South Africa

All five-herbivore species had a greater change in browse species utilisation after the addition of PEG when compared to standard game lick blocks, with a noteworthy effect on the diet of Burchell’s zebra and impala (Panel b). After the addition of PEG, the number of different browse species utilised in the diet of the common eland, impala, greater kudu and Burchell’s zebra increased. Before the deployment of the game lick blocks eland had similar diets in the study areas however after the addition of PEG the number of browse species utilised significantly increased identifying a notable shift in the diet of the eland exposed to PEG. The figure also shows that the changes in the diet after the application of the PEG game lick blocks are more significant than the changes after the deployment of the standard game lick blocks in all five-herbivore species (Panel b).

For the standard game lick blocks, there was no significant change in the percentage of browse diet of the impala (mixed feeder, eating both grass and browse). There was a slight change in favour of the utilisation of normally unpalatable browse species. After the introduction of the PEG game lick blocks, there was a large change in the percentages of browse species utilised and an increase in the utilisation of browse species that had low palatability.

The greater kudu and common eland had no major change in their diet of grass and browse with the addition of the standard game lick blocks. After the consumption of PEG there was a major increase in the utilisation of browse with higher secondary compounds. For blue wildebeest there was no substantial change in their diet between grass and browse neither with nor without the introduction of the standard game lick blocks or PEG game lick blocks (Figures 2 and 3, and Supporting Information D and E).

### Minerals in the faeces

There were minor comparative differences found in the mineral concentrations in the faeces of the five species of herbivore after utilising either the standard game lick block or the game lick block with the addition of PEG (Figure 4, Table 4). The average concentration of N found in the faeces of the common eland, blue wildebeest and Burchell’s zebra was higher after the utilization of the PEG game lick block, as was the average concentration of P in the faeces of the common eland, greater kudu and blue wildebeest, compared with the standard lick block (Supporting Information F). However, this difference was only statistically significant in the concentration of P in the faeces of the blue wildebeest. Impala was the only species that showed statistically significant higher levels of N and P after the utilisation of the standard game lick blocks over the PEG game lick blocks (Figure 4, Table 4). The concentration of Ca, S, Cu and B was significantly higher in the common eland after the utilisation of the PEG game lick blocks compared with the standard game lick blocks.

**Figure 4.**
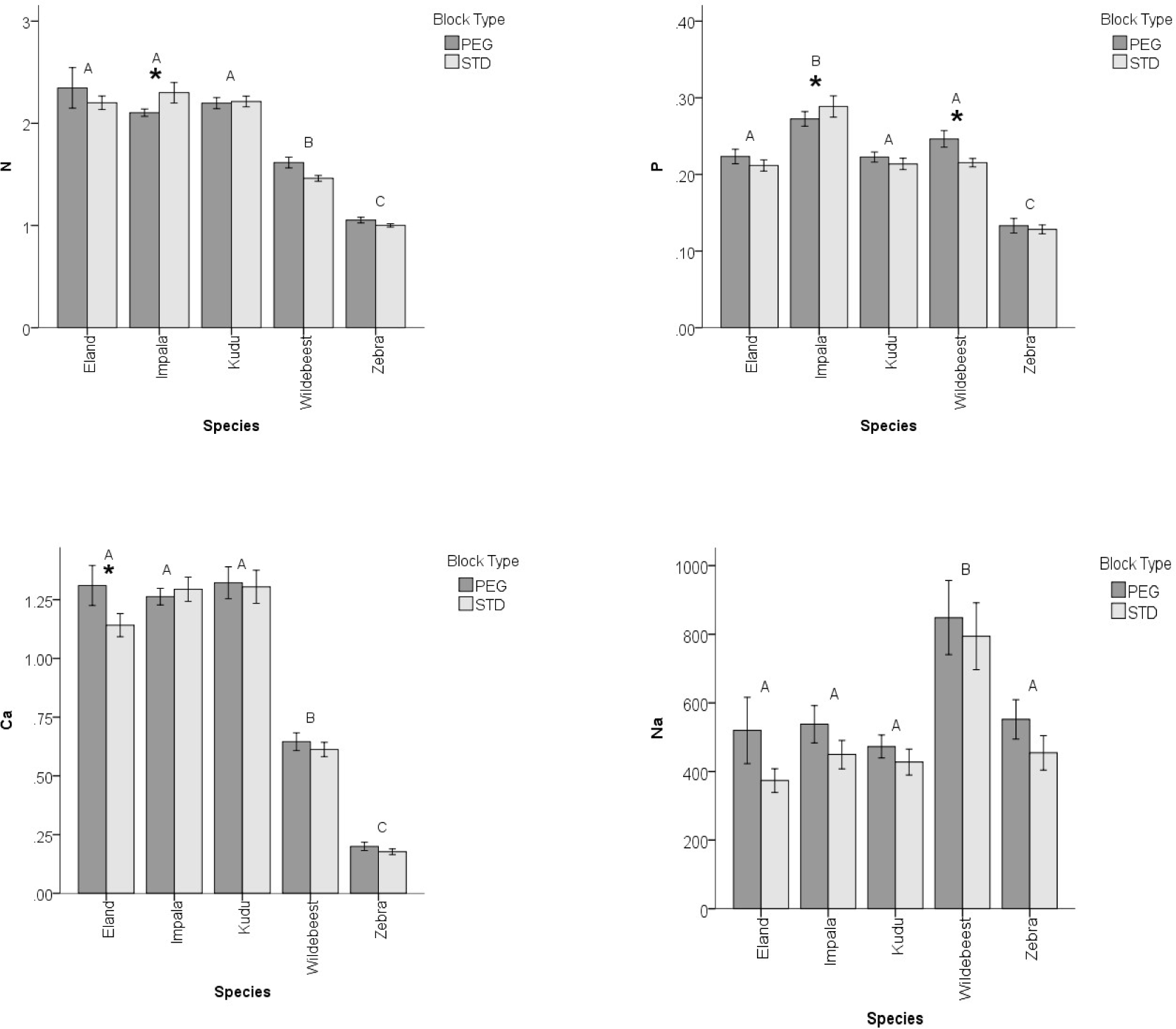
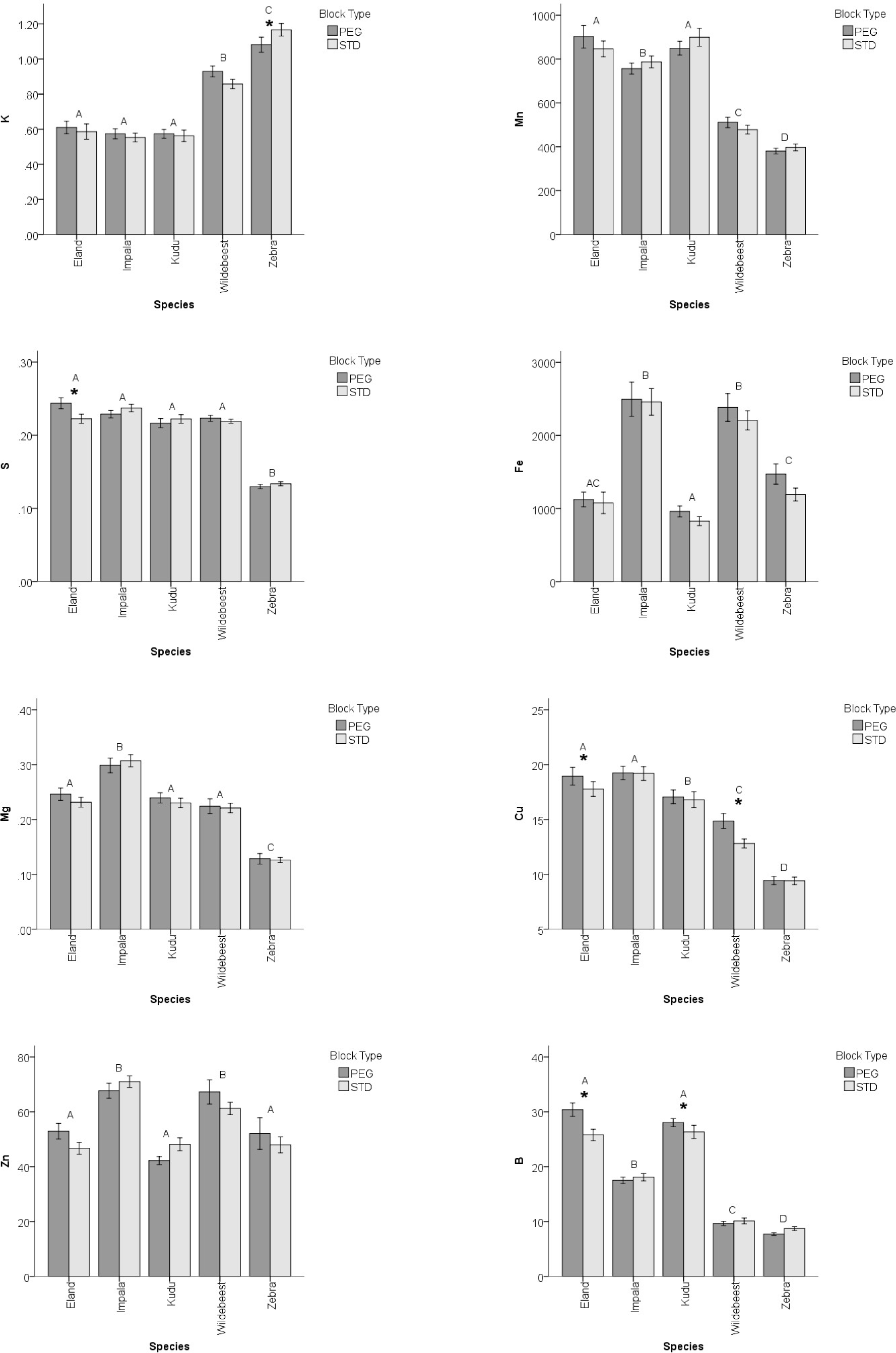
Mineral values in the faeces of the five-herbivore species (*viz*. impala (*Aepyceros melampus*), greater kudu (*Tragelaphus strepsiceros*), common eland (*Taurotragus oryx*), Burchell’s zebra (*Equus quagga burchellii*) and the blue wildebeest (*Connochaetes taurinus*) showing the effect of the availability of Polyethylene glycol (PEG) game lick blocks or Standard (STD) game lick blocks. The analysis of the minerals in the faeces samples were either recorded as a percentage as in the case of N, Ca, Mg, K, S and P or as mg/kg as with Na, Fe, Mn, Cu, Zn and B. Mineral values in the faeces are a measure of nutrient concentrations in the diet. Error bars represent the standard error of the mean. Letters indicate significant differences in concentrations of a mineral between the five herbivore species faeces and the star indicates the statistically significant differences in the mineral values of the faeces between the PEG and STD game lick blocks within the same herbivore species, in the treatments based on Šidák multiple comparisons tests using a linear mixed model (Table 4). The faecal samples were collected over a three-year period, 2016 – 2018, in the Welgevonden Game Reserve, South Africa

**Table 4.** Results of the linear mixed model for differences in minerals in the faeces between the Standard (STD) and Polyethylene glycol (PEG) game lick blocks (“block type”) for the five herbivore species (*viz*. impala (*Aepyceros melampus*), greater kudu (*Tragelaphus strepsiceros*), common eland (*Taurotragus oryx*), Burchell’s zebra (*Equus burchellii*) and the blue wildebeest (*Connochaetes taurinus*). Year and Site were the random factors, estimation methods were REML and the sample size n = 648 (see Figure 4 for the graphs). Akaike Information Criterion (AIC) estimates the prediction error of the amount of information lost in the model and establishes the quality of the model. This data was collected over a three-year period, 2016 – 2018, on the Welgevonden Game Reserve, South Africa. N is nitrogen, P is phosphorus, Ca is calcium, Na is sodium, and N.S. is not significant at the 5% level

|  |  | **N** | **P** | **Ca** | **Na** |
| --- | --- | --- | --- | --- | --- |
| Herbivore species | F | 163.9 | 85.0 | 273.9 | 12.2 |
|  | df | 634.3 | 634.2 | 631.2 | 633.4 |
|  | P | <0.001 | <0.001 | <0.001 | <0.001 |
| Block Type | F | 0.04 | 0.03 | 2.3 | 7.2 |
|  | df | 637.7 | 637.6 | 617.2 | 636.3 |
|  | P | N.S. | N.S. | N.S. | <0.01 |
| Herbivore species x Block Type | F | 3.4 | 2.3 | 1.0 | 0.1 |
|  | df | 634.5 | 634.8 | 633.0 | 634.3 |
|  | P | <0.01 | <0.01 | N.S. | N.S. |
| Year | Wald Z | 1.0 | 1.0 | 0.6 | 0.9 |
|  | P | N.S. | N.S. | N.S. | N.S. |
| Site | Wald Z | 0.7 | 0.9 | 1.3 | 1.0 |
|  | P | 0.507 | 0.388 | 0.197 | N.S. |
| AIC |  | 971.7 | -1491.5 | 541.4 | 9911.7 |
|  |  | **Mg** | **K** | **S** | **Fe** |
| Herbivore species | F | 92.7 | 121.2 | 193.1 | 49.8 |
|  | df | 634.6 | 634.2 | 633.2 | 634.6 |
|  | P | <0.001 | <0.001 | <0.001 | <0.001 |
| Block Type | F | 0.09 | 0.2 | 0.1 | 3.1 |
|  | df | 636.8 | 637.9 | 634.7 | 637.3 |
|  | P | N.S. | N.S. | N.S. | <0.10 |
| Herbivore species x Block Type | F | 0.7 | 2.1 | 2.2 | 0.3 |
|  | df | 635.1 | 634.4 | 633.6 | 635.2 |
|  | P | N.S. | <0.10 | <0.10 | N.S. |
| Year | Wald Z | 0.9 | 0.9 | 0.8 | 0.9 |
|  | P | N.S. | N.S. | N.S. | N.S. |
| Site | Wald Z | 1.2 | 1.6 | 1.1 | 1.1 |
|  | P | N.S. | N.S. | N.S. | N.S. |
| AIC |  | -1351.1 | 159.3 | -335.8 | 10883.7 |
|  |  | **Mn** | **Cu** | **Zn** | **B** |
| Herbivore species | F | 144.4 | 134.7 | 20.7 | 382.4 |
|  | df | 633.8 | 633.5 | 633.0 | 634.2 |
|  | P | <0.001 | <0.001 | <0.001 | <0.001 |
| Block Type | F | 0.08 | 1.3 | 0.4 | 7.2 |
|  | df | 635.2 | 635.3 | 626.4 | 636.6 |
|  | P | N.S. | N.S. | N.S. | <0.01 |
| Herbivore species x<br>Block Type | F | 1.2 | 2.3 | 1.3 | 3.7 |
|  | df | 634.0 | 633.2 | 634.7 | 635.0 |
|  | P | N.S. | <0.1 | N.S. | <0.01 |
| Year | Wald Z | 0.8 | 1.0 | 0.7 | 1.0 |
|  | P | N.S. | N.S. | N.S. | N.S. |
| Site | Wald Z | 1.3 | 0.3 | 0.9 | 1.1 |
|  | P | N.S. | N.S. | N.S. | N.S. |
| AIC |  | 8693.5 | 3657.9 | 6050.5 | 3946.8 |

## Discussion

Large herbivores in nutrient poor savannas have to deal with vegetation that generally has low nutrient concentrations, and hence low forage quality (Parker 2004; Mramba et al. 2018). These large herbivores, which are subjected to poor forage quality, coupled with high condensed tannins seem to have little access to large amounts and varieties of food, especially during the dry winter months, and thus are found to have low reproduction and survival rates in Welgevonden Game Reserve (Parker 2004; Nguyen et al. 2005; Hassanpour et al. 2011; Texeira et al. 2012; Mr. J Swart, 2020. Welgevonden Game Reserve, personal communication).

The hypothesis tested that deploying game lick blocks with PEG, compared to the standard game lick blocks, in a nutrient poor savanna will result in a broadening of herbivore diet, with higher percentages of browse species and increased utilisation per browse species that have high levels of condensed tannins and generally low to average palatability. With the addition of PEG, there was an increase in the utilisation of browse species in the diets of the common eland, impala, greater kudu and Burchell’s zebra on Welgevonden Game Reserve. The increase in the percentage of browse utilised was particularly significant in the diet of Burchell’s zebra and impala. The expectation was that the effect on the dietary choices would have had a greater impact on herbivore species that are browsers than on herbivore species that are grazers, as secondary compound level (tannins) in browse vegetation is generally higher than grasses (Barroso et al. 2001). However, we show that Burchell’s zebra (a grazer) and impala (a mixed feeder) have the largest change in diet selection after the addition of PEG, including a significant shift in vegetation choice from grass to browse. This change in diet due to PEG has potential further repercussions for the vegetation on Welgevonden Game Reserve due to the change in foraging behaviour. The implication is that larger amounts of browse could potentially be utilised, especially during the dry months than would normally be, resulting in a potential long-term decrease in closed woodlands to more open woodland system. Blue wildebeest (a grazer) had a decrease in the utilisation of browse after the addition of PEG in their diet, alluding to the fact that the effect of PEG may be species specific to the herbivore, specific to individual species dietary requirements or a combination of both.

With the addition of PEG, there is a general increase in the utilisation of vegetation with high tannin contents with medium and lower palatability ratings for the common eland, impala, greater kudu and Burchell’s zebra: the changes in the diets are positively correlated with the low and intermediate grazing rating and the low and average palatability of the individual plant species. It is shown in other studies on domestic animals that the addition of PEG to the diet does affect diet choice (Foley and Hume 1987; Marsh et al. 2003; Jansen et al. 2007; Mkhize et al. 2015; Mkhize et al. 2016), which collaborates with our findings for free-roaming herbivores in a nutrient poor savanna.

It can be deduced that PEG has a significant effect on the broadening of the diet choices of four of the five free-roaming herbivores on Welgevonden Game Reserve. The change in diet of these four free-roaming herbivores supports the findings of studies on penned animals that show that tannins act as both digestibility reducers and feeding deterrents that negatively affect intake (Mkhize et al. 2015) and for which PEG has been shown to reduce these adverse effects (Salawu et al. 1997; Landau et al. 2003). For blue wildebeest, classified as a grazer, there was no substantial change in their diet between graze and browse neither with nor without the introduction of the standard game lick blocks or PEG game lick blocks (Figures 2 and 3, and Supporting Information D and E).

We tested a further hypothesis to analyse if the change in diet, due to the addition of game lick blocks with and without the addition of PEG, had an effect on the concentration of minerals within the faeces of the five free-roaming herbivores. Despite the changes in diet after the utilisation of PEG, there was minimal variation between the levels of N and P in the faeces for the majority of herbivore species. The exceptions were the impala where N and P in the faeces were lower than the standard lick block after the utilisation of PEG, and for blue wildebeest which showed increases in faecal N and P after utilisation of PEG. Faecal nutrient content analyses are useful as short-term indicators of diet selection and nutrient status of free-roaming herbivores (Botha and Stock 2005; Codron et al. 2005). Grant et al. (2000) posit that faecal N levels can indicate a dietary deficiency which may precipitate nutritional stress, while faecal P levels indicate a deficiency that may lead to low reproductive rates in herbivores. Although the diet choices for common eland, greater kudu, impala and Burchell’s zebra broadened when PEG game lick blocks were introduced, there was no clear differences in the minerals in the faeces. Given the nutritional differences between the plant species, a possible explanation for these results is that the animals absorbed and utilised the extra nutrients consumed from their broader diet, which is necessary for survival in such nutrient poor environments (Parker 2004; Putman and Staines 2004; Texeira et al. 2012). Rainfall tends to reduce forage quality and the area becomes dominated by grass species with lower leaf N and P concentrations (Hopcraft et al. 2012). N and P concentrations in dry faeces of ruminants below the threshold of potential concern values of 1.2% and 0.25%, respectively (Prins 1996; Wrench et al. 1997; Grant et al. 2011), would constitute an exceedance of these thresholds and would be a concern for animal survival. Based on the N and P faeces threshold values of Wrench et al. (1997) and Grant et al. (2000), the Burchell’s zebra is the only animal with a concern pertaining to N in the study area, and the impala after utilising PEG is the only species with no concern pertaining to P (Supporting Information F). This shows that the addition of N and P to a nutrient poor system is imperative for the long-term survival of free-roaming herbivore species. Furthermore, when looking at the individual minerals identified in the faecal analysis in the five study animals, the two grazers (Burchell’s zebra and blue wildebeest), have the lowest amount of minerals available to them suggesting if the addition of minerals into a nutrient poor savanna is considered, the minerals should be applied to grass land areas which would be more accessible to the grazer species.

### Practical implications

Our study has shown that with the use of polyethylene glycol (PEG), four of the five free-roaming wild herbivores (*viz*. common eland, impala, greater kudu and Burchell’s zebra) in a nutrient poor savanna are able to utilise a greater number of otherwise less palatable tree species with higher levels of condensed tannins. This could allow these herbivore species to obtain more nutrients and energy from the nutrient poor diet, especially during the dry season, when many of the female herbivores in this dystrophic savanna are gravid. Alleviating the effects of food limitation by providing PEG may have a positive repercussion on fecundity levels. Our study concurs with the findings of Scogings et al. (2014) who stated that browsers and mixed feeders foraging preferences are strongly shaped by secondary compounds, however our findings confirms that this is also the case with Burchell’s zebra, which had a significant shift in diet from graze to browse. Our experiments thus potentially strengthen the ability of wild free-roaming wild herbivores to forge the critical link in the chain of reasoning between individual performance and individual food limitation on the one hand and population limitation and population dynamics on the other hand. This concurs with findings by Decandia et al. (2000) and Mkhize et al. (2015, 2016, 2018), who used PEG in feed trials with goats, where they concluded that PEG can improve the nutritive value of the feed and hence optimising the goat’s performance. These results lead to the conclusion that PEG is a valuable addition to the diet of the free-roaming wild herbivore species we studied.

These findings contribute to a better understanding of the diet of large herbivores in nutrient poor savannas and how management can act to improve foraging conditions, more especially in the dry months. In order to conclusively prove that PEG can break food limitation, additional information is required through further research to establish birth and survival rates, coupled with body condition change, of herbivores which have and have not utilised PEG in their diet.

## SUPPORTING INFORMATION

**Supporting Information A.** Standard Game Lick Block Composition (25kg blocks) as reported by the manufacturer WES Feeds (Thabazimbi, South Africa) (WES Feeds 2019)

| Nutrient | Minimum | Maximum | Unit |
| --- | --- | --- | --- |
| Protein | 150 |  | g/kg |
| Calcium | 15 |  | g/kg |
| Phosphorus | 8 |  | g/kg |
| Magnesium | 6 |  | g/kg |
| Sulphur | 6 |  | g/kg |
| Manganese | 200 |  | mg/kg |
| Copper |  | 40 | mg/kg |
| Cobalt |  | 0.5 | mg/kg |
| Iron |  | 100 | mg/kg |
| Iodine |  | 2 | mg/kg |
| Zinc |  | 350 | mg/kg |
| Selenium |  | 1 | mg/kg |
| Vitamin A | 20000 |  | iu/kg |
| Vitamin E | 30 |  | iu/kg |

**Supporting Information B.** Browse species located in the faeces with their digestibility ratings. The digestibility ratings were established through the use of the field guide to trees of Southern Africa (Van Wyk and Van Wyk 1997) and through the independent advice of experts in the field (Dr R. Grant (https://orcid.org/0000-0002-8419-5209), Dr M. Peel (https://orcid.org/0000-0003-1284-3665), Dr H. Bezuidenhout (http://orcid.org/0000-0002-6519-7321), Professor N. Owen-Smith (https://orcid.org/0000-0001-8429-1201), Professor K. Eloff (https://orcid.org/0000-0003-1494-9842), Mr J. Swart (Welgevonden Game Reserve Ecologist) and Mr G. Canning (https://orcid.org/0000-0001-6100-9541). The “X” in the table refers to the median when referring to the digestibility of these browse species for wild herbivores in the study area as described by the independent experts

| Browse Species | Infrequently consumed by a few intermediate feeders and browsers | Often consumed by some intermediate feeders and browsers | Very often consumed by many different intermediate feeders and browsers |
| --- | --- | --- | --- |
| *Senegalia burkei* |  |  | X |
| *Vachellia karoo* |  | X |  |
| *Apodytes dimidiata* | X |  |  |
| *Bauhinia petersiana* | X |  |  |
| *Berchemia zeyheri* |  |  | X |
| *Brachylaena rotundata* | X |  |  |
| *Burkea africana* | X |  |  |
| *Clerodendrum glabrum* | X |  |  |
| *Clutia pulchella* | X |  |  |
| *Combretum apiculatum* |  | X |  |
| *Combretum hereroense* |  | X |  |
| *Combretum nelsonii* | X |  |  |
| *Combretum zeyheri* |  | X |  |
| *Cryptolepis oblongifolia* | X |  |  |
| *Cussonia spicata* | X |  |  |
| *Dichapetalum cymosum* | X |  |  |
| *Dichrostachys cinerea* |  | X |  |
| *Diplorhynchus condylocarpon* |  | X |  |
| *Dombeya rotundifolia* | X |  |  |
| *Ehretia rigida* |  | X |  |
| *Elephantorrhiza burkei* | X |  |  |
| *Englerophytum magalismontanum* | X |  |  |
| *Erythrina lysistemon* | X |  |  |
| *Euclea crispa* | X |  |  |
| *Euclea linearis* | X |  |  |
| *Euclea natalensis* | X |  |  |
| *Faurea saligna* | X |  |  |
| *Flueggea virosa* |  | X |  |
| *Gardenia volkensii* |  |  | X |
| *Grewia bicolor* |  |  | X |
| *Grewia flavescens* |  | X |  |
| *Gymnosporia tenuispina* | X |  |  |
| *Indigofera melanadenia* | X |  |  |
| *Jasminum multipartitum* | X |  |  |
| *Lannea discolor* | X |  |  |
| *Lannea edulis* | X |  |  |
| *Lipkea javanica* |  | X |  |
| *Maytenus undata* | X |  |  |
| *Mundulea sericea* |  | X |  |
| *Ochna pulchra* | X |  |  |
| *Ozoroa paniculosa* | X |  |  |
| *Ozoroa sphaerocarpa* | X |  |  |
| *Parinari capensis* |  | X |  |
| *Pavetta gardeniifolia* | X |  |  |
| *Pellaea calomelanos* | X |  |  |
| *Peltophorum africanum* | X |  |  |
| *Protea caffra* | X |  |  |
| *Protea welwitschii* | X |  |  |
| *Pseudolachnostylis maprouneifolia* |  | X |  |
| *Psydrax livida* | X |  |  |
| *Pterolobium stellatum* | X |  |  |
| *Rhoicissus tridentata* | X |  |  |
| *Searsia pyroides* | X |  |  |
| *Spirostachys africana* |  | X |  |
| *Strychnos pungens* |  | X |  |
| *Syzygium cordatum* | X |  |  |
| *Syzygium guineense* | X |  |  |
| *Tarchonanthus camphoratus* |  | X |  |
| *Tarenna supra-axillaris* | X |  |  |
| *Terminalia brachystemma* | X |  |  |
| *Turraea obtusifolia* | X |  |  |
| *Vangueria infausta* |  | X |  |
| *Waltheria indica* | X |  |  |

**Supporting Information C.** Grass species found in the faeces with their grazing values obtained from the guide to grasses of Southern Africa (Van Oudtshoorn 2002). The “X” in the table refers to the median when referring to the digestibility of these grass species for wild herbivores in the study area as described by Van Oudtshoorn, 2002. ‘Grazing value’ is generally not known for wild herbivores with their different physiologies. The reported grazing values in this Table reflect more than a century of experiential knowledge form graziers in Southern Africa. Wildlife managers in Southern Africa have found these assessments generally useful for many of the wild grazers

| Grass Species | Grazing value low | Grazing value average | Grazing value high |
| --- | --- | --- | --- |
| *Andropogon chinensis* |  | X |  |
| *Aristida sp* | X |  |  |
| *Brachiaria brizantha* |  | X |  |
| *Brachiaria serrata* |  | X |  |
| *Cenchrus ciliaris* |  |  | X |
| *Chloris pycnothrix* | X |  |  |
| *Chloris virgata* |  | X |  |
| *Chrysopogon serrulatus* |  |  | X |
| *Cynodon dactylon* |  |  | X |
| *Dactyloctenium aegyptium* |  | X |  |
| *Digitaria eriantha* |  |  | X |
| *Diheteropogon amplexans* |  | X |  |
| *Eragrostis sp* |  | X |  |
| *Heteropogon contortus* |  | X |  |
| *Hyparrhenia filipendula* |  | X |  |
| *Hyparrhenia hirta* |  | X |  |
| *Hyperthelia dissoluta* |  | X |  |
| *Imperata cylindrica* | X |  |  |
| *Leersia hexandra* |  |  | X |
| *Loudetia simplex* |  | X |  |
| *Melinis nerviglumis* |  | X |  |
| *Melinis repens* | X |  |  |
| *Microchloa caffra* | X |  |  |
| *Monocymbium ceresiiforme* |  | X |  |
| *Panicum dregeanum* |  | X |  |
| *Panicum maximum* |  |  | X |
| *Panicum natalense* | X |  |  |
| *Paspalum scrobiculatum* |  | X |  |
| *Paspalum urvillei* |  | X |  |
| *Pogonarthria squarrosa* | X |  |  |
| *Sacciolepis typhura* | X |  |  |
| *Schizachyrium jeffreysii* | X |  |  |
| *Schmidtia pappophoroides* |  |  | X |
| *Setaria lindenberghiana* |  |  | X |
| *Setaria pumila* | X |  |  |
| *Setaria sphacelata* |  |  | X |
| *Sporobolus pyramidalis* | X |  |  |
| *Themeda triandra* |  | X |  |
| *Trachypogon spicatus* | X |  |  |
| *Triraphis schinzii* | X |  |  |
| *Tristachya biseriata* |  | X |  |
| *Urelytrum agropyroides* | X |  |  |
| *Urochloa mosambicensis* |  |  | X |

**Supporting Information D.**
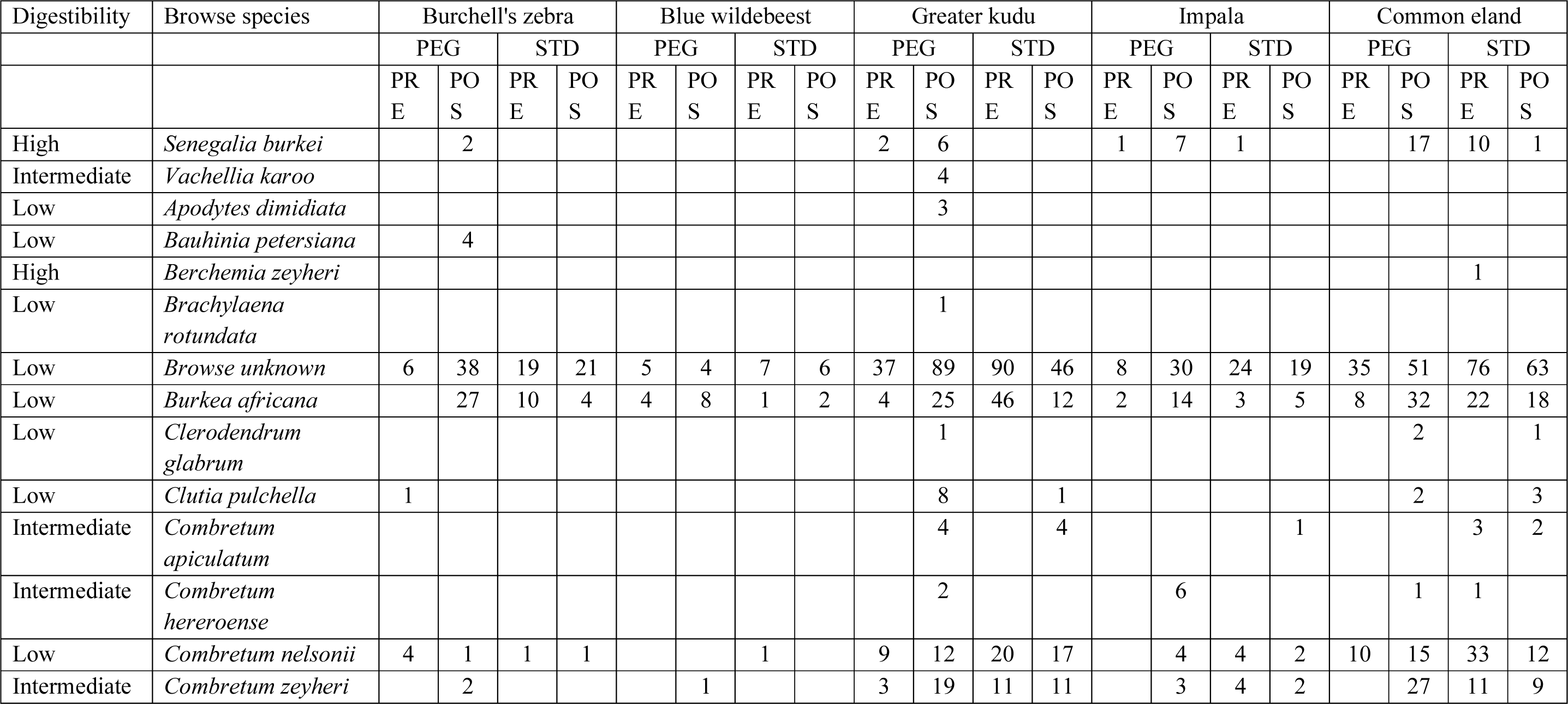
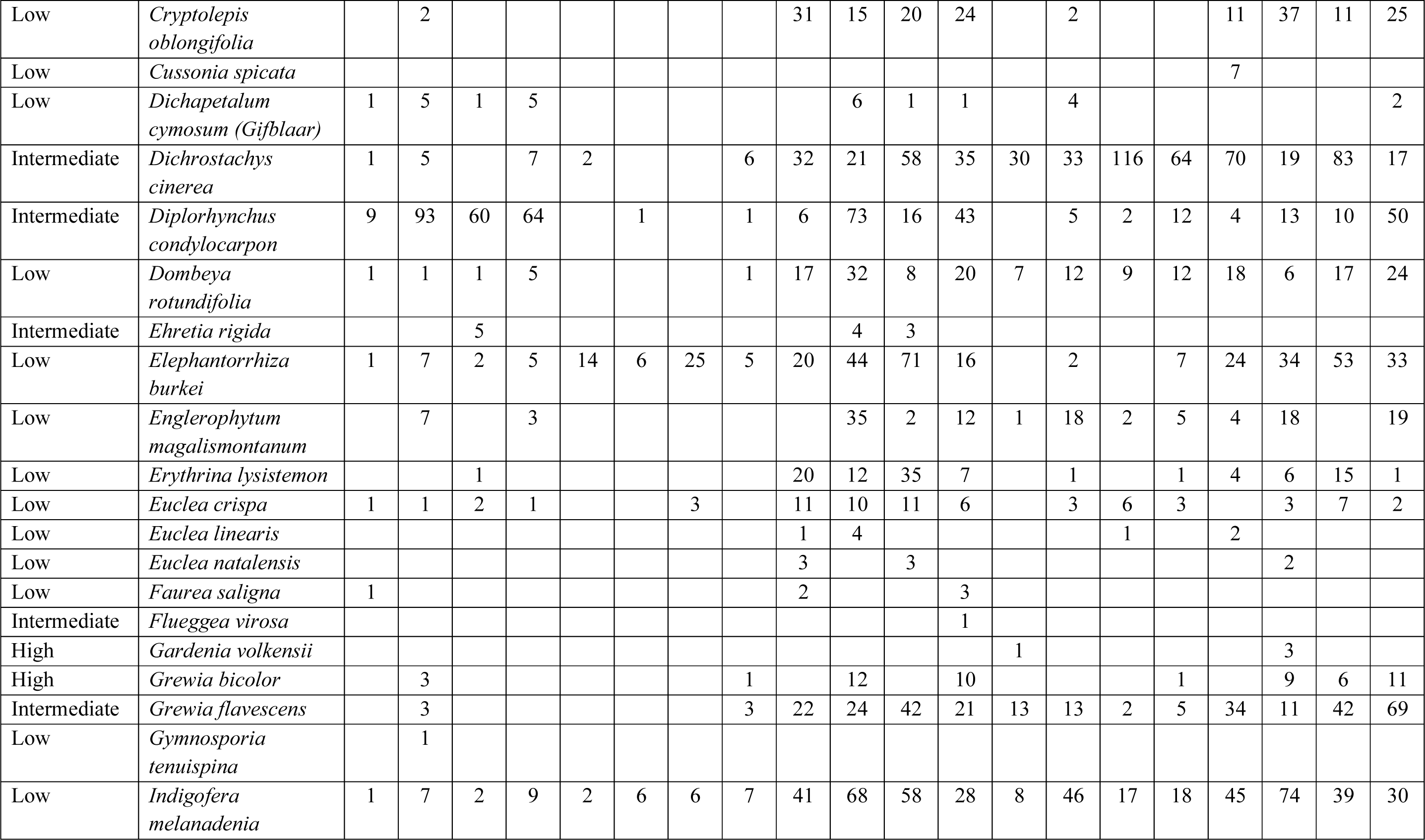
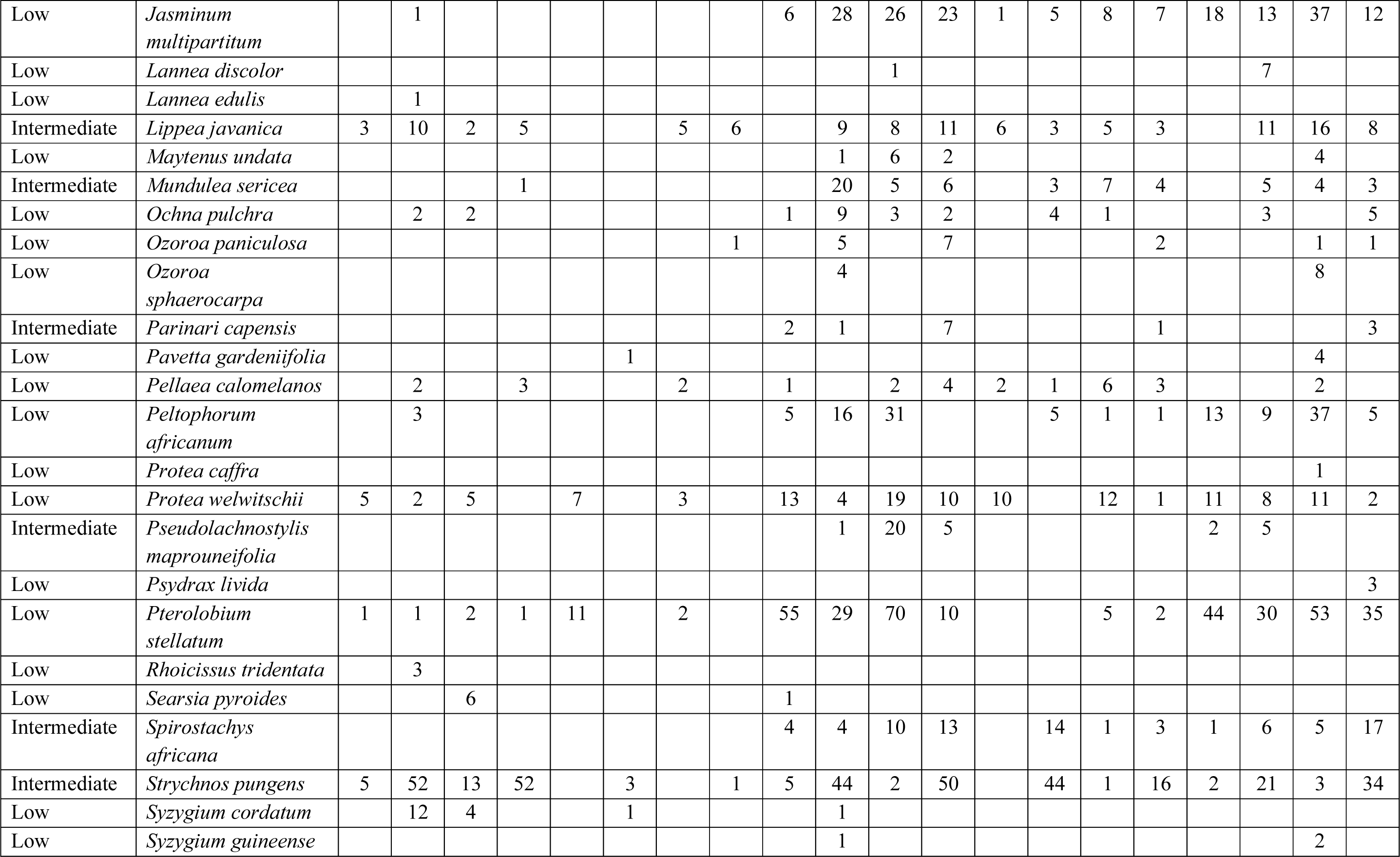
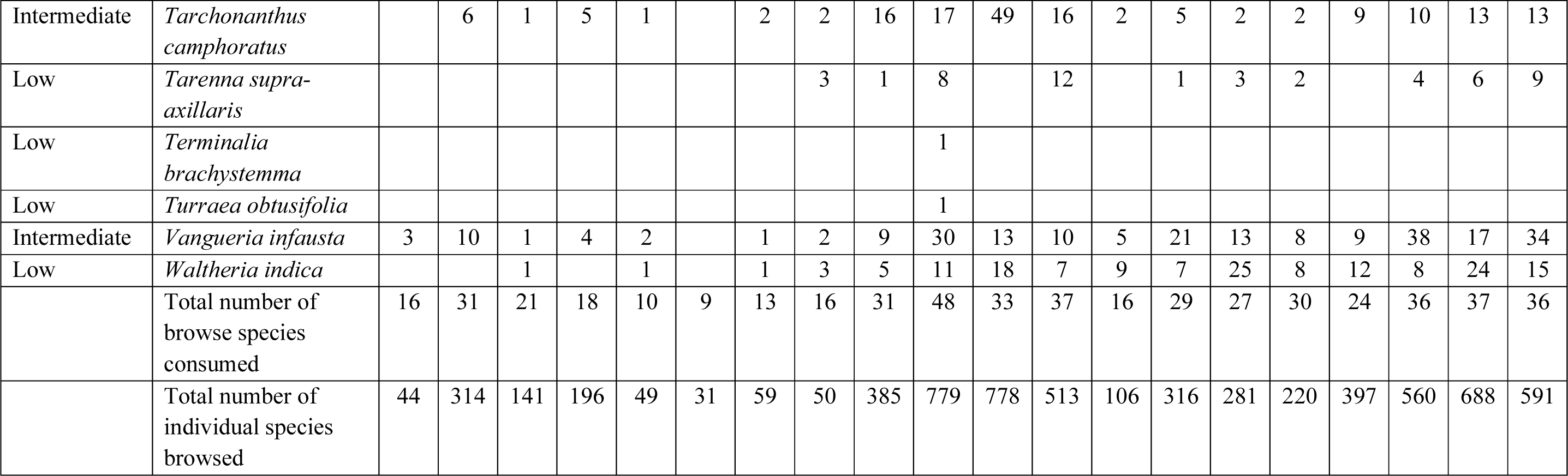
Diet change in browse after the utilization of game lick blocks with and without the supplementation of Polyethylene Glycol (PEG), for the five-herbivore species, impala (*Aepyceros melampus*), greater kudu (*Tragelaphus strepsiceros*), common eland (*Taurotragus oryx*), Burchell’s zebra (*Equus quagga burchellii*) and blue wildebeest (*Connochaetes taurinus*). In the Table, PEG refers to game lick blocks supplemented with Polyethylene Glycol. STD (for ‘standard’) refers to game lick blocks without PEG. PRE (for ‘prior’) refers to the analysis prior to the supplementation of PEG. POS (for ‘post’) refers to the analysis after the supplementation of PEG. The numbers in the Table refer to the number of each individual plant species found in the faeces of each animal species before or after the utilisation of PEG or STD lick blocks

**Supporting Information E.** Diet change in grass choice after the utilization of game lick blocks with and without the supplementation of Polyethylene Glycol, for the five-herbivore species, impala (*Aepyceros melampus*), greater kudu (*Tragelaphus strepsiceros*), common eland (*Taurotragus oryx*), Burchell’s zebra (*Equus quagga burchellii*) and the blue wildebeest (*Connochaetes taurinus*). In the Table, PEG refers to game lick blocks supplemented with Polyethylene Glycol. STD (for ‘standard’) refers to game lick blocks without PEG. PRE (for ‘prior’) refers to the analysis prior to the supplementation of PEG. POS (for ‘post’) refers to the analysis after the supplementation of PEG. The numbers in the Table refer to the number of each individual plant species found in the faeces of each animal species before or after the utilisation of PEG or STD lick blocks

| Grazing value | Graze species | Burchell's zebra |  |  |  | Blue wildebeest |  |  |  | Greater kudu |  |  |  | Impala |  |  |  | Common eland |  |  |  |
| --- | --- | --- | --- | --- | --- | --- | --- | --- | --- | --- | --- | --- | --- | --- | --- | --- | --- | --- | --- | --- | --- |
|  |  | PEG |  | STD |  | PEG |  | STD |  | PEG |  | STD |  | PEG |  | STD |  | PEG |  | STD |  |
|  |  | PRE | POS | PRE | POS | PRE | POS | PRE | POS | PRE | POS | PRE | POS | PRE | POS | PRE | POS | PRE | POS | PRE | POS |
| Average | *Andropogon chinensis* | 37 | 19 | 51 | 18 | 11 | 12 | 12 | 28 |  |  |  | 1 | 17 | 10 | 10 | 7 |  | 2 |  |  |
| Low | *Aristida sp* | 90 | 110 | 111 | 95 | 70 | 103 | 101 | 73 | 3 | 1 | 13 | 13 | 58 | 39 | 81 | 53 | 1 | 3 | 4 | 1 |
| Average | *Brachiaria brizantha* |  |  | 10 |  |  |  |  |  |  |  |  |  |  |  |  |  |  |  |  |  |
| Average | *Brachiaria serrata* | 20 | 3 | 6 | 8 | 1 | 13 | 6 | 7 |  |  |  | 2 | 13 | 3 | 7 | 5 |  |  |  |  |
| High | *Cenchrus ciliaris* | 1 | 1 | 1 | 1 | 3 | 2 | 1 |  |  |  |  |  |  |  |  |  |  |  |  |  |
| Low | *Chloris pycnothrix* | 12 | 9 | 6 | 7 |  | 3 | 20 | 7 |  |  | 1 |  | 8 | 5 | 8 | 4 |  | 5 |  | 1 |
| Average | *Chloris virgata* |  |  |  |  | 7 |  | 1 |  |  |  |  |  |  |  | 2 |  |  |  |  |  |
| High | *Chrysopogon serrulatus* | 1 |  | 1 |  | 1 | 3 | 5 |  |  |  |  |  | 5 |  | 1 |  |  |  |  |  |
| High | *Cynodon dactylon* | 42 | 55 | 91 | 25 | 51 | 102 | 81 | 63 |  | 1 | 20 | 13 | 59 | 61 | 70 | 46 | 1 | 4 |  | 3 |
| Average | *Dactyloctenium aegyptium* |  | 13 | 1 |  | 3 | 3 | 5 | 1 |  |  |  |  | 3 | 8 | 6 | 8 |  |  |  |  |
| High | *Digitaria eriantha* | 20 | 7 | 4 | 12 | 3 | 15 | 2 | 5 |  |  |  |  | 4 | 4 | 3 | 1 |  |  |  |  |
| Average | *Diheteropogon amplexans* | 5 | 2 | 1 |  | 2 |  |  |  |  |  |  |  | 2 |  |  |  |  |  |  |  |
| Average | *Eragrostis sp* | 148 | 196 | 247 | 147 | 189 | 323 | 409 | 224 | 1 | 2 | 34 | 29 | 186 | 185 | 232 | 181 |  | 11 | 3 | 1 |
| Low | *Grass unknown* | 36 | 37 | 52 | 21 | 26 | 36 | 47 | 29 | 1 |  | 6 | 4 | 18 | 22 | 41 | 18 |  | 1 | 1 |  |
| Average | *Heteropogon contortus* | 16 | 43 | 27 | 17 | 9 | 39 | 79 | 21 |  | 2 | 5 | 6 | 3 | 9 | 17 | 8 | 1 | 5 |  |  |
| Average | *Hyparrhenia filipendula* | 4 |  | 1 |  |  |  |  |  |  |  |  |  | 5 |  |  |  |  |  |  |  |
| Average | *Hyparrhenia hirta* |  |  |  | 1 |  |  |  |  |  |  |  |  |  |  |  |  |  |  |  |  |
| Average | *Hyperthelia dissoluta* | 1 |  |  |  |  |  |  |  |  |  |  |  | 1 |  |  |  |  |  |  |  |
| Low | *Imperata cylindrica* |  |  | 2 |  | 4 | 1 | 5 | 2 |  |  |  | 1 | 1 |  | 1 |  |  |  |  |  |
| High | *Leersia hexandra* |  | 2 | 3 | 3 |  | 7 | 5 | 9 |  |  |  | 5 | 1 |  | 2 | 4 |  | 1 |  |  |
| Average | *Loudetia simplex* | 1 |  | 1 | 1 | 1 | 1 |  |  |  |  |  |  |  |  |  |  |  |  |  |  |
| Average | *Melinis nerviglumis* |  |  | 2 | 1 |  |  | 1 |  |  |  | 2 |  |  | 1 | 2 |  |  |  |  |  |
| Low | *Melinis repens* | 22 | 24 | 29 | 23 | 23 | 31 | 24 | 29 |  | 1 | 5 | 5 | 25 | 9 | 30 | 12 |  | 3 |  | 1 |
| Low | *Microchloa caffra* |  |  |  |  |  | 1 | 2 |  |  |  | 1 |  | 2 |  |  |  |  |  |  |  |
| Average | *Monocymbium ceresiiforme* |  | 3 | 3 | 1 | 5 | 1 | 3 | 1 |  |  |  |  | 5 |  |  |  |  |  |  |  |
| Average | *Panicum dregeanum* |  |  | 2 |  |  |  |  |  |  |  |  |  |  |  |  |  |  |  |  |  |
| High | *Panicum maximum* | 27 | 8 | 33 | 6 | 4 | 15 | 21 | 21 |  |  | 1 | 3 | 22 | 3 | 28 | 6 |  | 2 |  |  |
| Low | *Panicum natalense* |  |  | 2 |  |  | 13 |  |  |  |  |  |  | 1 |  |  | 1 |  |  |  |  |
| Average | *Paspalum scrobiculatum* | 5 | 10 | 11 | 3 | 3 | 7 | 10 | 3 |  |  |  |  | 2 | 3 | 6 |  |  | 2 |  |  |
| Average | *Paspalum urvillei* |  | 1 |  |  |  | 2 |  | 1 |  |  |  | 2 |  | 1 |  |  |  |  |  |  |
| Low | *Pogonarthria squarrosa* |  |  |  |  |  |  |  |  |  |  |  |  | 1 |  |  |  |  |  |  |  |
| Low | *Sacciolepis typhura* |  | 3 | 1 |  | 1 | 3 | 2 | 2 |  |  |  |  |  |  | 1 |  |  |  |  |  |
| Low | *Schizachyrium jeffreysii* | 1 |  | 5 |  | 2 |  | 2 |  |  |  |  |  | 1 |  | 4 |  |  |  |  |  |
| High | *Schmidtia pappophoroides* | 8 | 6 | 14 | 7 | 3 | 9 | 21 | 7 |  |  | 15 |  | 8 | 1 | 38 | 3 |  |  | 1 | 1 |
| High | *Setaria lindenberiana* |  | 12 |  | 1 |  |  |  |  |  |  |  |  |  | 10 |  |  |  |  |  |  |
| Low | *Setaria pumila* | 1 |  |  |  |  |  |  |  |  |  |  |  |  |  |  | 2 |  |  |  |  |
| High | *Setaria sphacelata* | 8 | 8 | 7 | 2 | 2 | 9 | 1 | 6 |  |  |  |  | 6 | 2 | 7 | 3 |  |  |  |  |
| Low | *Sporobolus pyramidalis* | 2 |  |  |  |  |  |  |  |  |  |  |  | 1 |  |  |  |  |  |  |  |
| Average | *Themeda triandra* | 41 | 5 | 29 | 9 | 18 | 5 | 11 | 11 |  |  |  |  | 26 | 5 | 14 | 18 |  |  |  |  |
| Low | *Trachypogon spicatus* | 3 |  |  |  |  |  | 1 |  |  |  |  |  |  |  |  |  |  |  |  |  |
| Low | *Triraphis schinzii* |  |  | 1 | 1 | 2 | 1 | 3 |  |  |  |  |  |  |  | 2 |  |  |  |  |  |
| Average | *Tristachya biseriata* | 2 |  |  | 2 |  |  |  |  |  |  |  |  |  |  |  |  |  |  |  |  |
| Low | *Urelytrum agropyroides* |  |  |  |  |  | 1 |  |  |  |  |  |  |  |  |  |  |  |  |  |  |
| High | *Urochloa mosambicensis* | 2 |  | 2 |  | 3 |  |  |  |  |  | 1 |  | 3 |  | 2 |  |  |  |  |  |
|  | **Total number of grass species consumed** | **27** | **23** | **32** | **24** | **26** | **28** | **28** | **21** | **3** | **5** | **12** | **12** | **29** | **19** | **25** | **18** | **3** | **11** | **4** | **6** |
|  | **Total number of individual species grazed** | **583** | **577** | **757** | **412** | **447** | **761** | **881** | **550** | **5** | **7** | **104** | **84** | **487** | **381** | **615** | **380** | **3** | **39** | **9** | **8** |

**Supporting Information F.** Average percentage (with the 95% confidence intervals) of N- and P-concentrations found in the faeces for the five-herbivore species, impala (*Aepyceros melampus*), greater kudu (*Tragelaphus strepsiceros*), common eland (*Taurotragus oryx*), Burchell’s zebra (*Equus quagga burchellii*) and blue wildebeest (*Connochaetes taurinus*), following the use of game lick blocks with PEG (= PEG) or standard game lick blocks (= STD)

| **Herbivore species** | **Average N-PEG** | **Average N-STD** | **Average P-PEG** | **Average P-STD** |
| --- | --- | --- | --- | --- |
| Common eland | 2.35 ($\pm 0.40$) | 2.17 ($\pm 0.14$) | 0.22 ($\pm 0.02$) | 0.21 ($\pm 0.01$) |
| Impala | 2.10 ($\pm 0.07$) | 2.30 ($\pm 0.20$) | 0.27 ($\pm 0.02$) | 0.29 ($\pm 0.03$) |
| Greater kudu | 2.20 ($\pm 0.11$) | 2.21 ($\pm 0.10$) | 0.22 ($\pm 0.01$) | 0.21 ($\pm 0.01$) |
| Blue wildebeest | 1.62 ($\pm 0.10$) | 1.46 ($\pm 0.06$) | 0.25 ($\pm 0.02$) | 0.22 ($\pm 0.01$) |
| Burchell's zebra | 1.05 ($\pm 0.05$) | 1.00 ($\pm 0.03$) | 0.13 ($\pm 0.02$) | 0.13 ($\pm 0.01$) |

## Notes

### Competing Interest Statement

The authors have declared no competing interest.

